# A robust genome and assembly with transcriptomic data from the striped scorpion, *Centruroides vittatus*

**DOI:** 10.1101/2023.08.04.551372

**Authors:** Tsunemi Yamashita, Douglas D. Rhoads, Jeff Pummill

## Abstract

Scorpions, a seemingly primitive, stinging arthropod taxa, are known to exhibit marked diversity in their venom components. These venoms are known for their human pathology, but also important as models for therapeutic and drug development applications. We report a high quality genome assembly and annotation of the striped bark scorpion, *Centruroides vittatus*, created with several shotgun libraries. The final assembly is 760 Mb in size, with a BUSCO score of 97.8%, a 30.85% GC, and a N50 of 2.35 Mb. We estimated 36,189 proteins with 37.32% assigned to GO terms in our GOanna analysis. We were able to map 2011 and 60 venom toxin genes to contigs and scaffolds, respectively. We were also able to identify expression differences between venom gland (telson) and body tissue (carapace) with 19 Sodium toxin and 14 Potassium toxin genes to 18 contigs and two scaffolds. This assembly along with our transcriptomic data, provides further data to investigate scorpion venom genomics.

## Introduction

Scorpions are an ancient and diverse arthropod taxa primarily known for their medical importance and seemingly little morphological change over millions of years (Sharma et al. 2015, Lourenco 2018). Although all scorpions have a similar bauplan, they show immense variation in their venom components (Sunagar et al. 2013, Sharma et al. 2015, Housley et al. 2017, Lourenco 2018). The scorpion genus *Centruroides* constitutes the most medically important and one of the most diverse and wide-ranging scorpion taxa in North America (Gantenbein et al. 2001, Santibáñez-López et al. 2016, Esposito & Prendini 2019). The complex evolutionary history and geographic variability of this genus has generated controversy and taxonomic confusion (Sissom 1990, Borges et al. 2012, Santibáñez-López et al. 2016). The diversity of the venom also implies remarkable evolutionary adaptations with new and varied constituents discovered annually (Santibáñez-López et al. 2015, Housley et al. 2017). In spite of their medical importance, scorpion genomics has lagged behind venom transcriptomics and proteomics with only three of the estimated 2500 worldwide scorpion species with genome assembly entries in the NCBI database.

The scorpion *Centruroides vittatus* encompasses a large geographic range across the western USA and northern United Mexican States (Figure 1). Although a member of the toxic *Centruroides* genus, this species is not known as medically important (Kang & Brooks 2017). However, due to coevolution with mammalian predators, evidence suggests that western *C. vittatus* populations may possess a more medically significant venom than eastern populations (Rowe & Rowe 2008, Bowman et al. 2021). Throughout its geographic range, *C. vittatus* is commonly found in diverse ecological habitats, but populations across the northern and eastern geographic distributions appear to prefer dry, rocky south facing slopes or glade areas. Human introduction of this scorpion appears to also have created additional populations outside its known geographic range (Shelley & Sissom 1995). Here we present the assembled, annotated genome of *C. vittatus*. Integration of genome and transcriptome data show novel splicing and transcriptional activity around venom gene regions. Furthermore, this genome will complement the deposited genome of the more noxious western *C. sculpturatus*, and expand on analysis of this ancient taxon.

**Figure 1.**
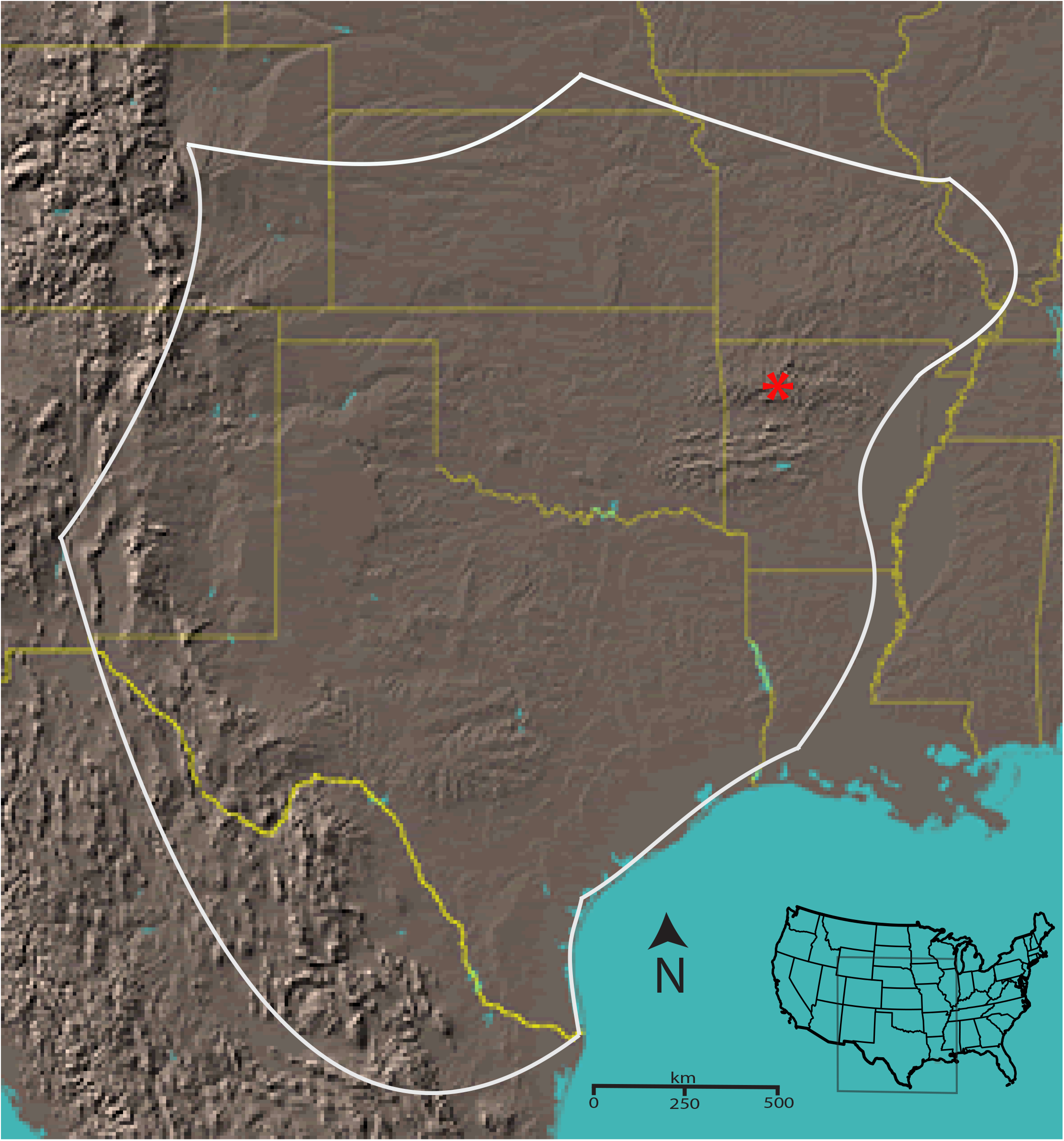
Approximate geographic range of *C. vittatus* with a red asterisk identifying the location for the individuals collected for the genomic analysis (from Yamashita et al. 2013).

## Materials and methods

### Genome sequencing and assembly

Total genomic DNA was extracted from four scorpions collected in Pope County, AR with the Qiagen genomic-tip and genomic DNA buffer set (Qiagen, Inc.). The genomic DNA quality and quantity was analyzed through 0.9% agarose gel electrophoresis, Qubit, and UV spectroscopy. One genomic DNA sample was sent to the University of Arkansas for Medical Sciences DNA Sequencing Core Facility for 300 base paired-end sequencing on a Illumina MiSeq. Two genomic DNA samples were sent to the National Center for Genome Resources (NCGR, NM) for PacBio 20K library generation and sequencing on 10 SMRT cell for each individual genome (Supp Table 1 & 2). A final sample was sent to the CTPR Genomics Lab at Arkansas Children’s Hospital Research Institute for a 300-cycle mid-output Illumina NextSeq genome sequencing.

The *de novo* assembly was conducted at the Arkansas High Performance Computing Center at the University of Arkansas. Sequence read data quality control check for Illumina short reads was conducted with FastQC (0.11.5; Andrews 2010) and trimmed with Trimmomatic (0.39; Bolger et al. 2014). For PacBio CLR long reads, NanoPlot (1.0.0; De Coster 2018) and FiltLong (0.2.0; Wick 2018) were utilized. The MiSeq reads were incorporated into the Flye assembly to correct the long PacBio reads. The two PacBio CLR long reads and the Illumina NextSeq quality trimmed reads were assembled with several software tools: MaSuRCA (V3.4.0; Zimin et al. 2013), Flye (V2.8.1; Kolmogorov et al. 2019), and also a version that was error corrected using Ratatosk (V0.1; Holley et al. 2020) and the Illumina data before assembly with Flye also utilizing consensus polishing via the tool incorporated with Flye. Draft assemblies were evaluated by two criteria: (i) the N50 statistic from contigs’ size, using QUAST v.5.0.2 (Gurevich et al., 2013), and (ii) the completeness score based on the presence of universal single copy ortholog genes, using BUSCO v.4.1.0 (Manni et al., 2021) against Arachnida ortholog dataset 10 (arachnida_odb10). Lastly, we identified and removed a unique *Mycoplasma* genome from our reads (Yamashita et al. 2019). The *Mycoplasma* genome was identified from the PacBio genomic sequence assembly as a unique 683,827 bp contig with a distinct GC content (43.7%) compared to the 30.85% GC content calculated for the scorpion contigs.

### Transcriptome assembly and annotation

Two male and one female scorpion were collected in northwest Arkansas, fed crickets with visual conformation of prey envenomation, then after three days, harvested for telson (venom gland) and carapace (body tissue) transcriptome analysis. The scorpions were flash frozen at −80 °C and total RNA extracted with a Trizol preparation (Sigma-Aldrich, St. Louis, MO, USA). RNA sample qc was analyzed through electrophoresis with an Aligent TapeStation system. RNA-seq with 50 bp reads was conducted at the University of Delaware on an Illumina genome sequencer (Illumina, Inc., San Diego, CA, USA). The data were viewed for initial quality through FastQC (v0.11.7), trimmed with Trimmomatic (v0.36; Bolger et al. 2014), and normalization of the data was performed using Trinity (v2.5.1; Haas et al. 2013). Assembly of the normalized reads was then performed with the following de novo assembly programs: Trinity (v2.5.1), SOAPdenovo2 (v2.4.1; Li et al. 2009), Velvet (v1.2.10; Zerbino et al. 2008), and TransAbyss (v1.5.4; Robertson et al. 2010) resulting in four individual assemblies. The transcriptome assemblies were then aggregated together using EviGene (Gilbert 2019), to remove redundancies, pics the best representatives, and filter out misassemblies. The final assembly was mapped to the genome assembly using NGen and quantified as RPKM using ArrayStar. RPKMs for contigs identified by BLASTn of the assembly for key genes were extracted for each sample over all contigs matching that BLAST query. In addition, the transcriptome assembly was blasted with a query database created from NCBI scorpion toxin and our current sodium toxin databases (2133 total toxin sequences). From these Blast searches, RPKM values for the two males and female were summarized for sodium toxin RNAs with additional searches for additional scorpion toxin RNAs. Transcriptome datasets were deposited in NCBI with the following IDs: TSA: GIPT01000000, SRA: SRR11917465, BioProject: PRJNA636371, BioSample: SAMN15075759

### Genome annotation

Repetitive elements were catalogued using RepeatModeler (V2.0.2a; Flynn et al. 2020) and repetitive elements masked with RepeatMasker (V4.1.2, Smit et al. 2013). The repeat masked genome was indexed and RNASeq reads from the carapace (body tissue) and the telson (venom gland) were aligned with STAR (V2.7.9a; Dobin et al. 2013) to create BAM files. Additionally, the RNASeq data was utilized to annotate the repeat masked scorpion genome with BRAKER (V2.1.6; Hoff et al. 2019). The predicted proteins from BRAKER were then utilized for a BLAST analysis of the TrEMBL database. Lastly, the polished *C*.*vittatus* genome with carapace and telson RNASeq BAM files, and the predicted, annotated polypeptides from BRAKER and the TrEMBL BLAST were loaded into IGV (V2.13.2a; Robinson et al. 2011) to view RNASeq Sashimi plots of the expression data relative to annotated exons. We also built a toxin BLASTn database with the polished *C. vittatus* genome against a sodium toxin query file housing 2133 sequences to further map toxin genes. The IGV visualizations to examine differential expression between carapace and telson were focused on contigs and scaffolds containing putative toxin genes based on the BLASTn toxin queries.

A functional annotation analysis used the Cyverse pipeline developed for arthropods (Saha et al. 2021). This pipeline combines outputs from GOanna (GO annotations) and InterProScan (functional motifs) as well as mapping proteins to pathways via KOBAS.

## Results and discussion

### Genome assembly and annotation

The four genome sequencing outputs resulted in three final *de novo C vittatus* genome assemblies (Table 1). Of the three final genome assemblies, the Ratatosk-Flye assembly was judged as the most complete as it exhibited the largest reduction in contig number, with the largest contig size and N50 (Table 1, Table S1). This assembly also showed the best BUSCO statistics with 97.8% complete (92.6% unique, 5.2% duplicates) (Table 2). The transcriptome basic statistics for the venom gland are presented in Table 3 with the repetitive element data in Table S2. The transcriptomic data and the annotated polypeptide data were incorporated into IGV to visualize gene expression variation between the venom gland and body tissue (Figures 2 – 4). The functional annotation workflow predicted 36,189 proteins, which is higher than the 17,364 proteins predicted in *L. hesperus* (Western Black Widow Spider) (Saha et al. 2021), but comparable to the *C. sculpturatus* protein number of 35,529. The GOanna and InterProScan results between *C. vittatus* and *L. hesperus* are comparable (Table 4.). The KOBAS output shows a marked difference with *C. vittatus* exhibiting higher numbers of proteins assigned to pathways and percent assigned to KEGG pathways when compared to *L. hesperus* (Table 5).

**Table 1:** Assembly statistics from three assembled genomes of *C. vittatus* and *C. sculpturatus*.

| Quast Assembly Statistics | MaSuRCA | Flye | Ratatosk-Flye | *C. sculpturatus* |
| --- | --- | --- | --- | --- |
| # contigs ($\geq 0$ bp) | 6383 | 2842 | 2071 | |
| # contigs ($\geq 50000$ bp) | 5787 | 2177 | 1694 | |
| Total length ($\geq 0$ bp) | 791,048,345 | 769,382,264 | 760,848,696 | 925,474,958 |
| # contigs | 6135 | 2440 | 1837 | 35,614 |
| Largest contig | 2,437,683 | 10,350,238 | 11,744,082 |  |
| Total length | 790,486,062 | 768,752,645 | 760,461,457 | 863,133,193 |
| GC (%) | 30.89 | 30.85 | 30.85 | 31.4 |
| N50 | 336,247 | 1,876,222 | 2,358,559 |  |

\*\*\* The MaSuRCA assembly incorporates the aggregated PacBio CLR long reads and the Illumina NextSeq reads. The Flye assembly only incorporates the aggregated PacBio CLR long reads, and the Ratatosk-Flye assembly incorporates the aggregated PacBio CLR long reads with Ratatosk correction with MiSeq short reads.

**Table 2:** Busco statistics of the three *C. vittatus* assembled genomes.

| Busco Statistics | MaSuRCA | % | Flye | % | Ratatosk-Flye | % |
| --- | --- | --- | --- | --- | --- | --- |
| Complete BUSCOs (C) | 2794 | 95.2 | 2842 | 96.8 | 2870 | 97.8 |
| Complete and single-copy BUSCOs (S) | 2615 | 89.1 | 2686 | 91.5 | 2716 | 92.6 |
| Complete and duplicated-BUSCOs (D) | 179 | 6.1 | 156 | 5.3 | 154 | 5.2 |
| Fragmented BUSCOs (F) | 23 | 0.8 | 26 | 0.9 | 14 | 0.5 |
| Missing BUSCOs (M) | 117 | 4.0 | 66 | 2.3 | 50 | 1.7 |
| Total Busco Groups searched | 2934 |  | 2934 |  | 2934 |  |

\*\*\* arachnida\_odb10.2019-11-20 lineage markers

**Table 3:** Transcriptome basic statistics from a combined *C. vittatus* telson (toxin gland) of three scorpions. From Bowman et al. 2021.

|  |  |
| --- | --- |
| Transcript number | 226,162 |
| Transcripts > 500 bp | 48,463 |
| Transcripts > 1000 bp | 26,909 |
| Average length of assembled transcripts (bp) | 463.164 |
| Longest transcript (bp) | 9,430 |
| Total length (bp) | 104,750,193 |
| Transcript N50 | 1,149 |

**Table 4:** Functional annotation analyses with Goanna and Interproscan results.

|  |  |  | GOanna<br>(Blast) |  | InterProScan<br>(Motif<br>Analysis) |  |
| --- | --- | --- | --- | --- | --- | --- |
| species | Proteins | Proteins<br>Assigned<br>GO terms | Proteins<br>Assigned<br>GO | Average<br>GAQ | Proteins<br>Assigned<br>GO | Average<br>GAQ |
| *Centruroides vittatus* | 36, 189 | 37.32% | 7.37% | ----- | 30.45% | ----- |
| *Latrodectus hesperus* | 17,364 | 31.17% | 2.02% | 197.44 | 30.44% | 28.896 |

\*\*\*Note: the *C. sculpturatus* protein number is estimated as 35,529.

**Table 5:**
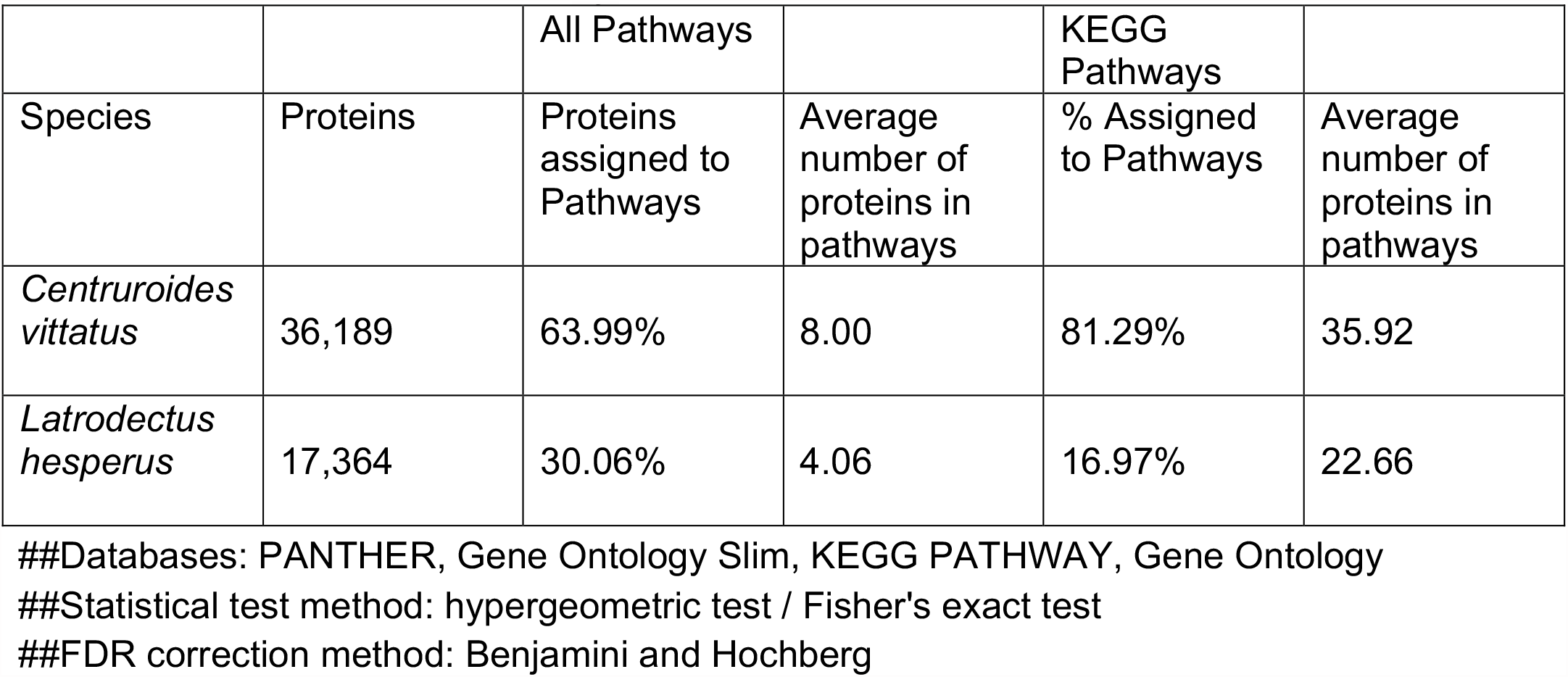
Functional annotation analyses with KOBAS results.

**Figure 2a.**
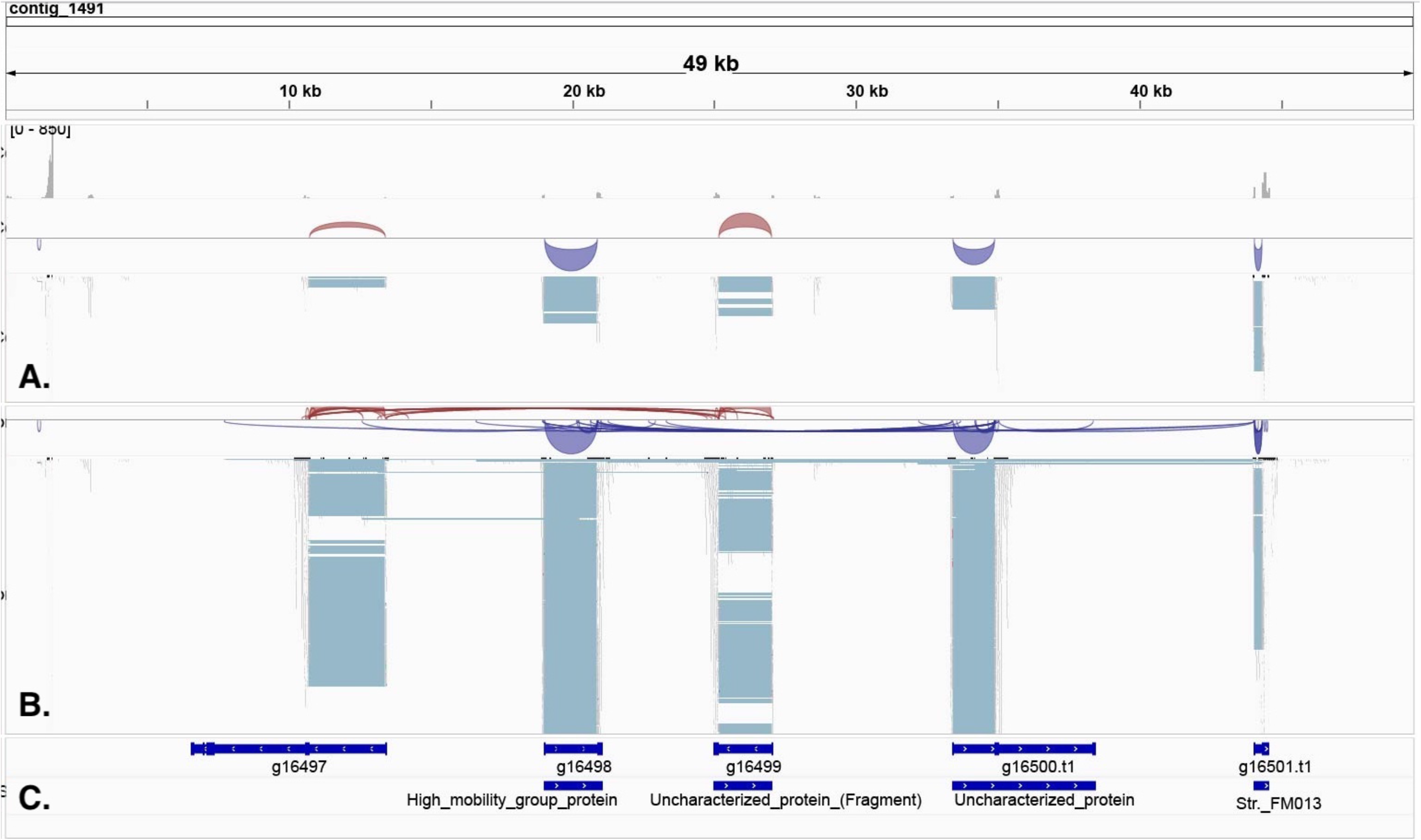
IGV visualization of contig 1491 (49Kb) showing putative sodium toxin gene expression mapped with respect to carapace (Panel A) and telson (Panel B). Predicted Proteins from BRAKER and the TrEMBL BLASTp results are mapped in Panel C. Gene expression variation is denoted adjacent to the pale blue blocks and suggests clustering of sodium toxin genes in this contig. The red and blue arcs in panels A & B represent splice junctions connecting exons. The red represents mRNA reads mapped to the + strand: The blue represents mRNA reads mapped to the – strand. The section of the contig that shows the five putative sodium toxin gene regions spans 35Kb with intervening genome sequence of 3 – 8Kb. The scales for mRNA read coverage are the same in panels A & B and range from 0 - 250 reads.

**Figure 2b.**
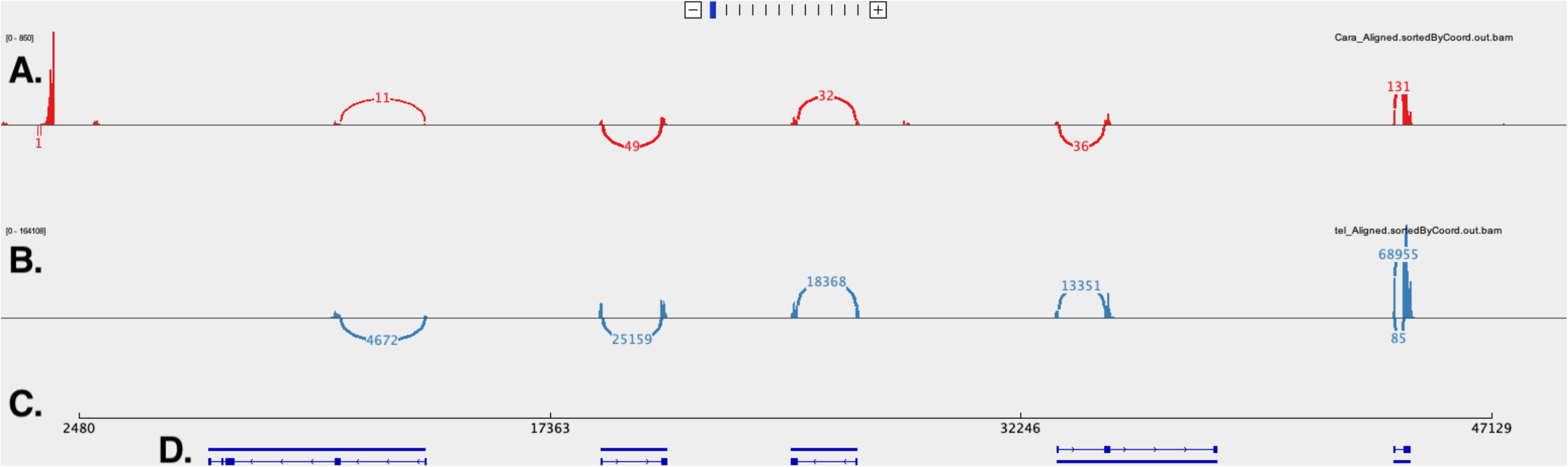
IGV visualization of sashimi plot for contig 1491 (49Kb) of the region shown in Fig 2A illustrating putative sodium toxin gene expression mapped with respect to carapace (A) and telson (B). The contig location is delineated in C. Predicted Proteins from BRAKER are mapped in D. Gene expression variation between the two tissues is denoted with thin red or blue bar graphs along A or B. Splice junctions are noted with red (carapace) or blue (telson) arcs connecting putative exons with the read number shown in the arcs and suggests clustering of sodium toxin genes in this contig. Arcs above the alignment track represent reads mapped to the + strand & arcs below the alignment track represent reads mapped to the – strand. The read numbers for the carapace (red) from left to right are 11, 49, 32, 36, and 131. The reads mapped to the telson (blue) from left to right are 4672, 25159, 18368, 13351, and 68955. The section of the contig that shows the five putative sodium toxin gene regions spans 35Kb with intervening genome sequence of 3 – 8Kb.

### Toxin gene specifics

The BRAKER annotation of the Ratatosk – Flye assembly mapped putative toxin genes to 2011 contigs and 60 scaffolds (Table 6). The mapping of toxin genes to the assembly with the scorpion toxin Blastn file refined the genes to a subset of 848 contigs and 57 scaffolds. Further analysis of the Blastn output identified putative toxin genes in 18 contigs and 2 scaffolds in which the toxin genes showed much higher expression in the telson (venom gland) vs. the carapace, including 19 putative sodium and 14 putative potassium toxins genes.

**Table 6:** Scorpion toxin gene expression mapped to the *C. vittatus* genomic assembly. The Ratatosk-Flye assembly totals represent the subset of contigs and scaffolds with identified putative toxin genes from BRAKER. The scorpion toxin Blastn file represents contigs and scaffolds with toxin gene hits from a query file of 2133 scorpion toxin genes. The total sodium and potassium toxin genes with higher telson expression mapped to the 18 contigs and 2 scaffolds from the filtered Blastn data are 19 & 14 genes, respectively.

|  | Toxin genes and percentages mapped to contigs | Toxin genes and percentages mapped to Scaffolds |
| --- | --- | --- |
| Ratatosk-Flye Assembly Totals | 2011 | 60 |
| Scorpion Toxin Blastn file | 846 (42%) | 57 (95%) |
| Filtered Scorpion Blastn Na/K results with higher telson expression than carapace | 17 Na toxins & 14 K toxins to 18 contigs | 2 Na Toxins to two scaffolds |

The toxin gene mapping suggests many toxin genes are only differentially expressed in body tissue versus venom glands, rather than uniquely expressed in venom glands. One contig that spans 35Kb (contig 1491) shows five paralogs of a putative sodium toxin gene in tandem, separated by intervening genomic sequences of 3 – 8Kb, suggesting ancestral gene duplication in this region (Fig. 2a). The IGV view of contig 1491 also suggests the sodium toxin genes in this region are arranged on both + and – DNA strands, which may indicate gene inversions. The Sashimi plot of this region (Fig. 2b.) clarifies the RNAseq mapping, showing markedly higher expression in telson mRNAs in this putative sodium toxin gene region.

Other contigs showed a pattern of larger genomic regions with mapped sodium toxin genes. For example, contig 2703 shows a 1900 bp region with putative sodium toxin genes mapped, suggesting multiple sodium toxin genes in this region (Fig 3.). Putative potassium toxin genes were located on other contigs and scaffolds, six versus 15 for putative sodium toxin genes, with no evidence of duplicated regions (e.g., Fig 4.). These genes also appear to exhibit differential expression in the telson vs the carapace rather than unique expression in the telson. These initial findings support a model of recent toxin gene duplication events that may underline the incredible sodium toxin diversity in the New world *Centruroides* species. (Rendon-Anaya et al. 2012, Drukewitz & von Reumont 2019).

**Figure 3.**
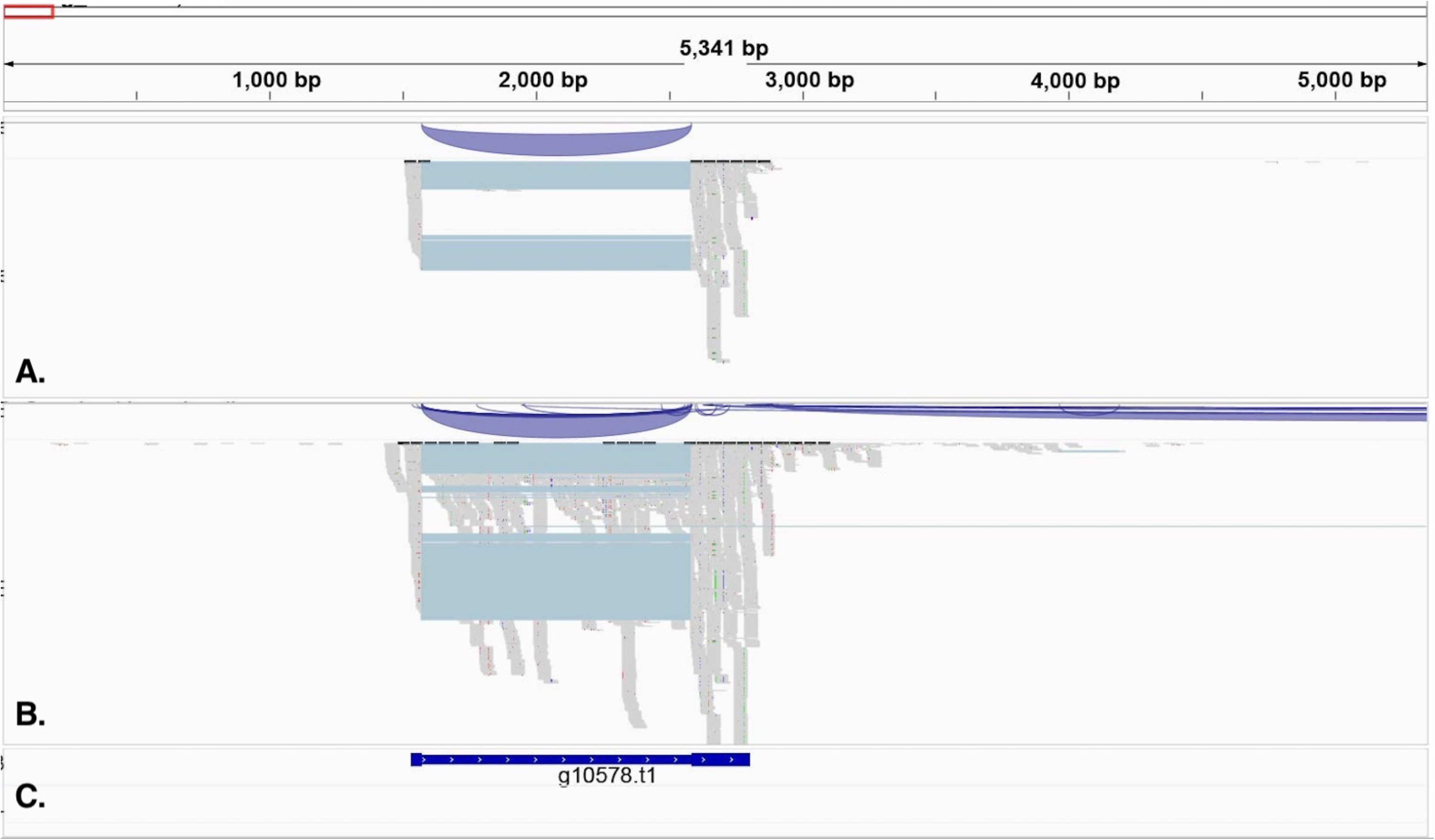
IGV visualization of a 5Kb region in contig 2703 (157Kb) showing putative sodium toxin gene expression mapped with respect to carapace (Panel A) and telson (Panel B). Predicted Proteins from BRAKER and the TrEMBL BLASTp results are mapped in Panel C. Gene expression variation is denoted with the 50bp reads shown in the grey regions and suggests clustering of sodium toxin genes in this contig. The section of the contig that shows the putative sodium toxin gene region spans 2150bp. The scales for mRNA read coverage are the same in panels A & B and range from 0 - 250 reads.

**Figure 4.**
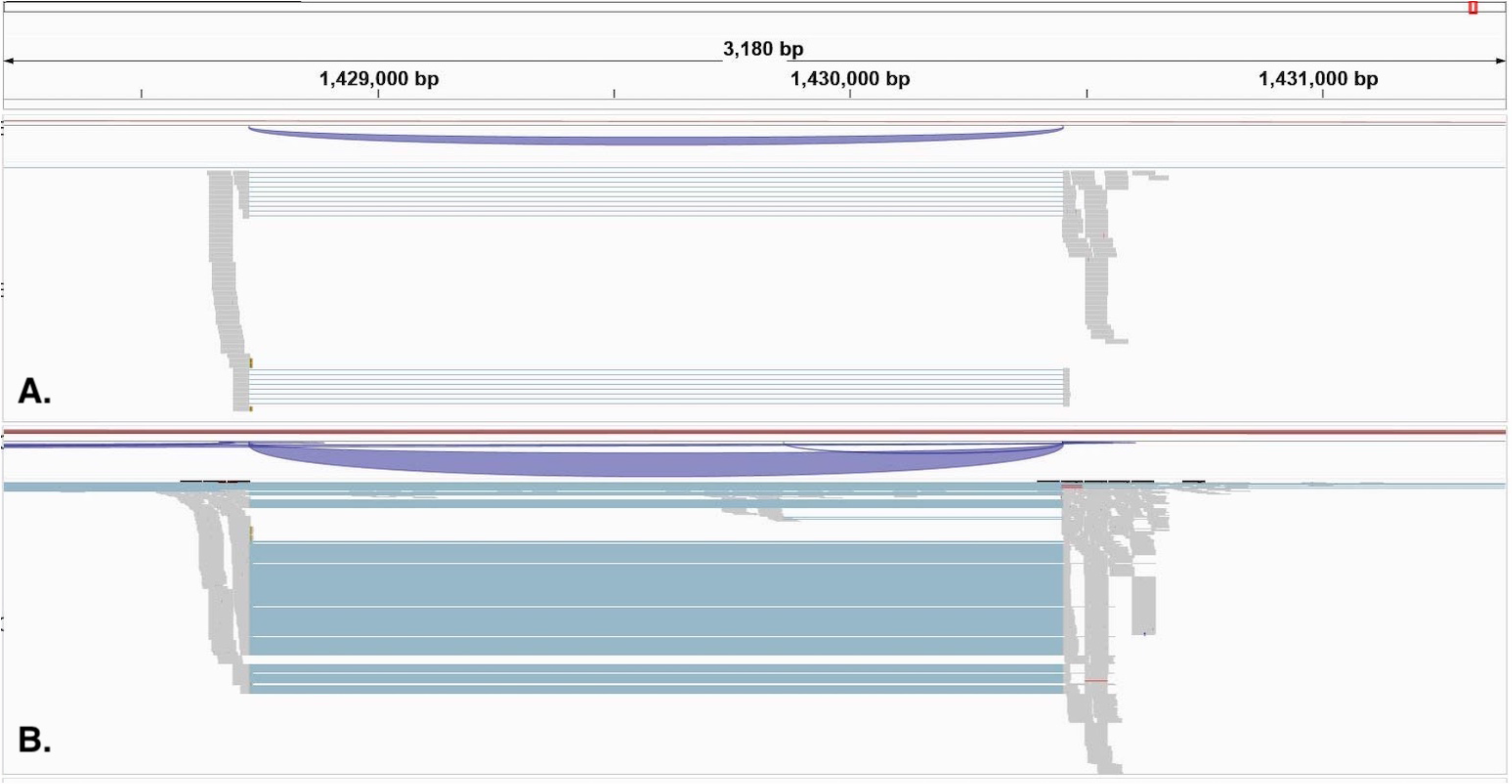
IGV visualization of a 2,300bp region in contig 82 (1,461Kb) showing putative potassium toxin (ERG1) gene expression mapped with respect to carapace (Panel A) and telson (Panel B). Gene expression variation between the two tissues is denoted with the grey regions with the 50 bp reads and the pale blue region is a putative intron gap of 1700bp between mapped reads. The scales for mRNA read coverage are the same in panels A & B and range from 0 - 250 reads.

## Conclusion

We describe a genomic assembly and annotation for the scorpion *Centruroides vittatus* coupled with transcriptomic data mapped to contigs and scaffolds. Our assembly shows a genome of 760Mb in length, with 98% of sequences mapped to 2071 contigs. Our results also highlight the substantial toxin gene diversity in this scorpion and show toxin gene expression patterns between body tissue and the venom gland. This genome will complement the growing number of venomous species with genomes in published databases.

## Data availability

The genome assembly was deposited at NCBI under accession number JASCZU000000000; BioProject PRJNA937744; BioSample SAMN33417986.

Supplementary material is available at G3 online.

## Acknowledgements

The authors thank Shane Sanders for conducting the RepeatModeler, RepeatMasker, STAR, and BRAKER analyses.

## Funding

Funding was provided by the Arkansas INBRE program, with a grant from the

National Institute of General Medical Sciences, (NIGMS), P20 GM103429 from the National Institutes of Health, USA.

## Literature cited

Andrews S. FastQC: A Quality Control Tool for High Throughput Sequence Data [Online].2010. Available online at: http://www.bioinformatics.babraham.ac.uk/projects/fastqc/

Bolger AM, Lohse M, Usadel B. Trimmomatic: A flexible trimmer for Illumina Sequence Data. Bioinformatics. 2014; 30(15): 2114–2120.

Borges A, Miranda RJ, Pascale JM. Scorpionism in Central America, with special reference to the case of Panama. J. Ven. Ani. Tox. Trop. Dis. 2012; 18: 130–143.

Bowman A, Fitzgerald C, Pummill JF, Rhoads DD, Yamashita T. Reduced Toxicity of Centruroides vittatus (Say, 1821) May Result from Lowered Sodium β Toxin Gene Expression and Toxin Protein Production. Toxins. 2021; 13(11):828.

Darling AC, Mau B, Blattner FR, Perna NT. Mauve: multiple alignment of conserved genomic sequence with rearrangements. Genome research. 2004; 14(7): 1394–1403.

De Coster W, D’Hert S, Schultz DT, Cruts M, Van Broeckhoven C. NanoPack: visualizing and processing long-read sequencing data. Bioinformatics. 2018;34(15):2666–2669.

Dobin A, Davis CA, Schlesinger F, Drenkow J, Zaleski C, Jha S, Batut P, Chaisson M, Gingeras TR. STAR: ultrafast universal RNA-seq aligner. Bioinformatics. 2013; 29(1): 15– 21.

Esposito LA, Prendini L. Island Ancestors and New World Biogeography: A Case Study from the Scorpions (Buthidae: Centruroidinae) [published correction appears in Sci Rep. 2020 Apr 30;10(1):7545]. Sci Rep. 2019;9(1):3500. Published 2019 Mar 5.

Evolution stings: the origin and diversification of scorpion toxin peptide scaffolds. Toxins (Basel). 2013;5(12):2456–2487. Published 2013 Dec 13.

Flynn JM, Hubley R, Goubert C, Rosen J, Clark AG, Feschotte C, Smit AF. RepeatModeler2 for automated genomic discovery of transposable element families. Proceedings of the National Academy of Sciences of the United States of America. 2020; 117(17): 9451–9457.

Gantenbein B, Fet V, Barker MD. Mitochondrial DNA reveals a deep, divergent phylogeny in Centruroides exilicauda (Wood, 1863) (Scorpiones: Buthidae). In: Fet V, Seldon PA, editors. Scorpions 2001. In Memoriam Gary A. Polis. British Arachnology Society: Burnham Beeches, Bucks, UK; 2001. p. 235–244.

Gilbert DG. Genes of the pig, Sus scrofa, reconstructed with EvidentialGene. PeerJ. 2019;7:e6374. Published 2019 Feb 1.

Gurevich A, Saveliev V, Vyahhi N, Tesler G. QUAST: quality assessment tool for genome assemblies. Bioinformatics. 2013; 29(8): 1072–1075.

Haas BJ, Papanicolaou A, Yassour M, Grabherr M, Blood PD, Bowden J, Couger MB, Eccles D, Li B, Lieber M, MacManes MD. et al. De novo transcript sequence reconstruction from RNA-seq using the Trinity platform for reference generation and analysis. Nat Protoc. 2013;8(8):1494–1512.

Hoff KJ, Lomsadze A, Borodovsky M, Stanke M. Whole-Genome Annotation with BRAKER. Methods in molecular biology. 2019; 1962: 65–95.

Holley G, Beyter D, Ingimundardottir H, Møller PL, Kristmundsdottir S, Eggertsson HP, Halldorsson BV. Ratatosk: hybrid error correction of long reads enables accurate variant calling and assembly. Genome Biol. 2021; 22(1): 28

Housley DM, Housley GD, Liddell MJ, Jennings EA. Scorpion toxin peptide action at the ion channel subunit level. Neuropharmacology. 2017;127:46–78.

Kang AM, Brooks DE. Nationwide Scorpion Exposures Reported to US Poison Control Centers from 2005 to 2015. J. Med. Toxicol. 2017; 13: 158–165.

Kolmogorov M, Yuan J, Lin Y, Pevzner PA. Assembly of long, error-prone reads using repeat graphs. Nat Biotechnol. 2019;37(5):540–546. doi:10.1038/s41587-019-0072-8

Koren S, Walenz BP, Berlin K, Miller JR, Bergman NH, Phillippy AM. Canu: scalable and accurate long-read assembly via adaptive k-mer weighting and repeat separation. Genome Res. 2017;27(5):722–736. doi:10.1101/gr.215087.116

Li R, Yu C, Li Y, Lam TW, Yiu SM, Kristiansen K, Wang J. SOAP2: an improved ultrafast tool for short read alignment. Bioinformatics. 2009;25(15):1966–1967.

Lourenço WR. The evolution and distribution of noxious species of scorpions (Arachnida: Scorpiones). J Venom Anim Toxins Incl Trop Dis. 2018;24:1. Published 2018 Jan 4.

Manni M, Berkeley MR, Seppey M, Zdobnov EM. BUSCO: Assessing Genomic Data Quality and Beyond. Curr Protoc. 2021;1(12):e323.

NCGAS/de-novo-transcriptome-assembly-pipeline (Version 4.0.1). Available online: Zenodo. http://doi.org/10.5281/zenodo.3945505 (accessed on 15 July 2019).

Robertson G, Schein J, Chiu R, Corbett R, Field M, Jackman SD, Mungall K, Lee S, Okada HM, Qian JQ, et al. De novo assembly and analysis of RNA-seq data. Nat Methods. 2010;7(11):909–912. doi:10.1038/nmeth.1517

Robertson G, Schein J, Chiu R, Corbett R, Field M, Jackman SD, Mungall K, Lee S, Okada HM, Qian JQ, Griffith M, et al. De novo assembly and analysis of RNA-seq data. Nat Methods. 2010;7(11):909–912.

Robinson JT, Thorvaldsdóttir H, Winckler W, Guttman M, Lander ES, Getz G, Mesirov JP. Integrative genomics viewer. Nat Biotechnol. 2011;29(1):24–26. doi:10.1038/nbt.1754

Rowe AH, Rowe MP. Physiological resistance of grasshopper mice (Onychomys spp.) to Arizona bark scorpion (Centruroides exilicauda) venom. Toxicon. 2008; 52: 597–605.

Saha S, Cooksey AM, Childers AK, Poelchau MF, McCarthy F. Workflows for rapid functional annotation of diverse arthropod genomes. Insects 2021: 12(8): 748.

Santibáñez-López CE, Francke OF, Ureta C, Possani LD. Scorpions from Mexico: From species diversity to venom complexity. Toxins. 2016; 8(1) 2.

Schulz MH, Zerbino DR, Vingron M, Birney E. Oases: robust de novo RNA-seq assembly across the dynamic range of expression levels. Bioinformatics. 2012;28(8):1086–1092.

Sharma PP, Fernández R, Esposito LA, González-Santillán E, Monod L. Phylogenomic resolution of scorpions reveals multilevel discordance with morphological phylogenetic signal. Proc Biol Sci.

Shelley RM, Sissom WD. Distributions of the scorpions Centruroides vittatus (say) and Centruroides hentzi (banks) in the United States and Mexico (scorpiones, buthidae). J. Arach. 1995; 23: 100–110.

Shi Y, Shang J. Long Noncoding RNA Expression Profiling Using Arraystar LncRNA Microarrays. Methods Mol Biol. 2016;1402:43–61.

Simão FA, Waterhouse RM, Ioannidis P, Kriventseva EV, Zdobnov EM. BUSCO: assessing genome assembly and annotation completeness with single-copy orthologs. Bioinformatics. 2015;31(19):3210–3212. doi:10.1093/bioinformatics/btv351

Sissom WD. Life history. In: The Biology of the Scorpions, Polis GA, editor. Stanford (CA) Stanford University Press, 1990. p. 161–223.

Smit, AFA, Hubley, R & Green, P. RepeatMasker Open-4.0. 2013-2015 < http://www.repeatmasker.org>.

Stothard P, Wishart DS. Circular genome visualization and exploration using CGView. Bioinformatics. 2005;21(4):537–539.

Sunagar K, Undheim EA, Chan AH, Koludarov I, Muñoz-Gómez SA, Antunes A, Fry BG. Teruel R, Fet V, Graham MR. The first mitochondrial DNA phylogeny of Cuban Buthidae (Scorpiones: Buthoidea). Boletín Soc. Entomológica Aragonesa. 2006; 39: 219−226.

Walker BJ, Abeel T, Shea T, Priest M, Abouelliel A. et al. Pilon: an integrated tool for comprehensive microbial variant detection and genome assembly improvement. PLoS One. 2014;9(11):e112963.

Yamashita T, Rhoads DR. Species delimitation and morphological divergence in the scorpion Centruroides vittatus (Say, 1821): Insights from phylogeography. PLoS ONE. 2013; 8(7): e68282

Yamashita T, Rhoads DD, Pummill J. Genome Analyses of a New Mycoplasma Species from the Scorpion Centruroides vittatus. G3 (Bethesda). 2019;9(4):993-997. Published 2019 Apr 9.

Zerbino DR, Birney E. Velvet: algorithms for de novo short read assembly using de Bruijn graphs. Genome Res. 2008;18(5):821–829.

Zimin AV, Marçais G, Puiu D, Roberts M, Salzberg SL, Yorke JA. The MaSuRCA genome assembler. Bioinformatics. 2013 Nov 1;29(21):2669–77.

