## Supplemental information for "A robust genome and assembly with transcriptomic data from the striped scorpion, *Centruroides vittatus*"

**Supplementary Information**

Table S1. Initial genome sequencing results from PacBio genome sequencing.

| Job Metric | Q1133 | Q1171 |
|---|---|---|
| Number of Bases | 6,812,411,387 | 9,669,253,726 |
| Number of Reads | 555,183 | 642,400 |
| N50 Read Length | 18,378 | 21,548 |
| Mean Read Length | 12,270 | 15,051 |
| Read Quality Score | 0.85 | 0.85 |

Table S2. Repetitive elements in the *C. vittatus* genome identified from a RepeatMasker output.

| Identity | Number of elements | Length occupied (bp) | Percentage of sequence (%) |
|---|---|---|---|
| Retro elements | 68,527 | 46,749,745 | 6.14 |
| Penelope | 28,475 | 12,237,284 | 1.61 |
| LINEs | 59,660 | 37,732,270 | 4.96 |
| R1/LOA/Jockey | 10869 | 10,675,340 | 1.4 |
| LTR elements | 8,867 | 9,017,475 | 1.19 |
| DNA transposons | 131,005 | 52,169,258 | 6.86 |
| Tc1-IS630-Pogo | 69,372 | 25,562,168 | 3.36 |
| Unclassified | 931,189 | 214,632,422 | 28.21 |
| Total Interspersed repeats | | 313,551,425 | 41.21 |
| Simple repeats | 268,925 | 11,706,345 | 1.54 |

***Only elements above 1% are reported.
